# Combinatorial Intracellular Delivery Screening (CIDS) of Anticancer Drugs

**DOI:** 10.1101/2020.08.28.271429

**Authors:** Belén Sola-Barrado, Diana Moreira Leite, Edoardo Scarpa, Aroa Duro-Castano, Giuseppe Battaglia

## Abstract

Conventional drug solubilisation strategies limit the understanding of the full potential of poorly water-soluble drugs during drug screening. Here, we propose a screening approach in which poorly water-soluble drugs are entrapped in poly (2-(methacryloyloxyethyl phosphorylcholine)-poly(2-(diisopropylaminoethyl methacryate) (PMPC-PDPA) polymersomes (POs) to enhance drug solubility and facilitate intracellular delivery. By using a human paediatric glioma cell model, we demonstrated that PMPC-PDPA POs mediated intracellular delivery of cytotoxic and epigenetic drugs by receptor-mediated endocytosis. Additionally, when delivered in combination, drug-loaded PMPC-PDPA POs triggered both an enhanced drug efficacy and synergy compared to that of a conventional combinatorial screening. Hence, our comprehensive synergy analysis illustrates that our screening methodology, in which PMPC-PDPA POs are used for intracellular co-delivery of drugs, allows us to identify potent synergistic profiles of anticancer drugs.

---

For any given medicine to be successful, the drug and its formulation need to be properly balanced. However, solubilisation, administration route and bioavailability are elements equally important too. Yet, they are often left to the clinical evaluation stage, deciding the fate of its trial. Formulation science, nowadays almost re-branded as nanomedicine, involves ever sophisticated tools to make sure that the active ingredient, the drug, reaches its target avoiding unwanted effects. Yet at the drug discovery process, conventional strategies for drug solubilisation limit the understanding of the full potential of poorly water-soluble molecules during drug screening.^1^ When testing the efficacy of a drug, the standard procedure commonly involves solubilisation of the drug in a suitable solvent followed by administration to *in vitro*/*in vivo* models. Three major challenges arise here. First, given that more than 40% of new chemical entities (NCEs) present a poor water-solubility,^2^ the use of organic solvents is usually necessary for their solubilisation, with consequent toxicity. Second, a poor water-solubility results in limited and variable drug bioavailability mainly due to a deficient absorption from the gastrointestinal tract to the blood.^3^ Last, the majority of drugs acts on intracellular targets, and thus an impaired intracellular delivery affects the efficacy of the drug.^4,5^ As we progress in personalised medicine, single drug therapy is not longer enough to ensure the desired therapeutic impact, instead, combination therapy, which consists on the use of two or more drugs in the same treatment, may result in a higher efficacy due to the pharmacological interaction of the different drugs. Combination therapy can also benefit from intracellular delivery, as a pharmacological interaction may be facilitated by granting access of the combined agents to their target sites. To this respect, nanomedicine can improve the drug screening process by providing an augmenting of drug solubility and intracellular delivery. ^6–9^

In the last decades, we have been working on the design and optimisation of a successful intracellular delivery system. We use poly (2-(methacryloyloxyethyl phosphorylcholine) - poly (2-(diisopropyl aminoethyl methacrylate) (PMPC-PDPA) polymersomes (POs), synthetic vesicles formed by self-assembly of amphiphilic copolymers. These synthetic vesicles are able of encapsulating and delivering a large variety of molecules including nucleic acids,^10,11^ proteins^12^ and poorly water-soluble drugs.^13^ PMPC-PDPA POs are internalised by binding to scavenger receptor class B type 1 (SR-B1), cluster of differentiation 81 (CD81) and cluster of differentiation 36 (CD36) receptors. ^14,15^ They thus deliver payloads intracellularly,^10,16,17^ escaping endosomal sorting via a prompt pH driven disassembly. ^18,19^ Here, we propose a screening approach for both single and combined cytotoxic and epigenetic drugs where PMPC-PDPA POs are used as a tool for drug solubilisation and intracellular delivery of a combinatory panel of anti-cancer drugs (Fig.1). We apply this approach to diffuse midline glioma H3K27M mutant (DMG-H3K27M) as a disease model. DMG-H3K27M is an aggressive diffuse paediatric astrocytoma with a life expectancy is *<* 2 years in more than 90% of the cases,^20–22^ which shows high molecular heterogeneity. ^22^ Initially, we optimised PMPC-PDPA POs for the intracellular delivery of the cytotoxic drugs paclitaxel (PTX, a vinca alkaloid) and carfilzomib (CRF, a proteosome inhibitor) and the epigenetic drug panobinostat (PNB, a histone dehacetylase) to DMG-H3K27M glioma cells. We then investigated the intracellular delivery of both single drugs and dual-delivery using PMPC-PDPA POs, and using the R package SynergyFinder,^23^ we provide the screening of synergistic interactions on a combinatory panel of anti-cancer drugs.

**Figure 1:**
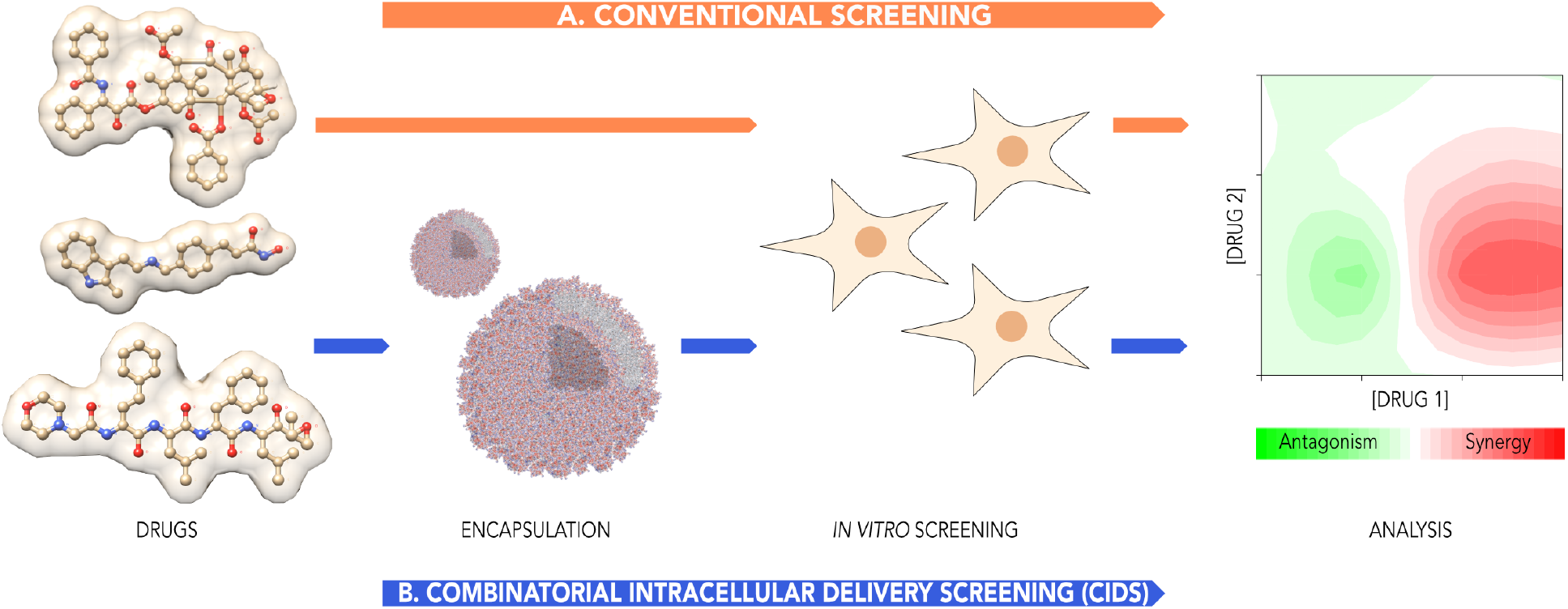
Combinatorial Intracellular Delivery Screening (CIDS) of anticancer drugs. (A) Conventional drug screening, in which the drugs are dissolved in organic solvents and then directly tested in cancer cells, and (B) combinatorial intracellular delivery screening (CIDS), in which drugs are encapsulated in PMPC-PDPA POs to facilitate solubilisation and mediate intracellular delivery in cancer cells. Following in vitro screening, synergy analysis is carried out using the R package SynergyFinder. Drugs tested are from top to bottom PXT, PNB and CRF.

We started by preparing PXT, CRF and PNB-loaded *PMPC*_25_ − *PDPA*_70_ POs by solvent switch method,^25^ and further purified and characterised by Transmission Electron Microscopy (TEM) and Dynamic Light Scattering (DLS). TEM showed that the block copolymers self-assembled into spherical vesicles (Fig.S1 A-D), which is in agreement with literature.^26^ As depicted in the inset in Fig.S1D, the membrane boundary of the PMPC-PDPA POs was identified with the phosphotungstic acid (PTA) negative staining, showing a uniform membrane thickness of ≈ 6 nm. Additionally, the size and polydispersity were confirmed by DLS (Table S1 and Fig.S2). Formulations of unloaded and drug-loaded POs exhibited a narrow size distribution with a hydrodynamic size of ≈ 70 nm and low polydispersity index (PDI < 0.2, except for PNB-POs which still showed a PDI < 0.5). Drug concentration, POs production efficiency and loading efficiency of the PTX-, CRF- and PNB-loaded POs were then determined by reverse-phase HPLC (Table S1). The three drugs were successfully encapsulated with a PO production efficiency (P.E.) from 5.90 to 30.25 % and showing loading efficiencies (L.E.) that seems to increase for higher partition coefficient (logP) values, suggesting that more hydrophobic drugs where more easily encapsulated.

In order to understand the interaction between PMPC-PDPA POs and paediatric glioma cells for the intracellular delivery of drugs, we selected three glioma cells lines: SF8628 (human paediatric, H3K27M mutation), cell line 7 (mouse, H3K27M mutation) and F98 (rat, undifferentiated glioma). Firstly, the expression of H3K27M mutation and SR-B1 receptors was evaluated by western blot analysis (Fig.S3A,B). H3K27M mutation and SR-B1 receptor expression was confirmed on SF8628 and Line 7, while F98 showed the absence of the H3K27M mutation and low levels of SR-B1. Thus, F98 was used as a negative control for our cellular studies. Subsequently, the three cells lines were incubated with unloaded PMPC-PDPA POs for 24 hours showing a cell viability > 75% for all the tested concentrations. These results indicate that PMPC-PDPA POs are safe to use as a tool for the drug screening in these glioma cells (Fig.S3C). Finally, kinetics of uptake of fluorescently-labelled POs in SF8628, line 7 and F98 cells were studied by confocal microscopy (Fig.S3D). Our results demonstrate that the PMPC-PDPA POs are internalised within 5 minutes in Line 7 and SF8628, while internalisation in F98 was only observed at 24 hours of incubation (Fig.S4). These cellular uptake profiles indicate that PMPC-PDPA POs are possibly internalised via SRB1-mediated endocytosis in SF8628 and Line 7. Considering the relatively fast uptake of POs and the presence of the genetic mutation of interest, we focused the remainder of our studies on human SF8628 cells as a reliable *in vitro* cell model for DMG-H3K27M.

Aiming to obtain a screening of the single drugs in the paediatric cell model, we tested the effect of PTX, CRF and PNB both free (solubilised in DMSO) and loaded within PMPC-PDPA POs. These drugs are currently approved by the FDA for clinical use in humans and have already shown their potential against glioma *in vitro*.^27–29^ These features, combined with their distinct mechanisms of action, make them great candidates for combinatorial screening approach using POs. We then tested the effect of three drugs individually both free and loaded in PMPC-PDPA POs (Fig.2 and S5-7). We observed that the treatment with drug-loaded POs led to a reduced cell viability compared to the free drugs. Particularly, the maximum efficacy of the treatment, expressed as cell viability, was: (1) for PXT 43.92 ± 12.97 % in loaded-POs compared to 71.20 ± 22.48 % free; (2) for CRF was 6.55 ± 0.44 % in loaded-POs compared to 10.68 ± 5.72 % free; (3) for PNB was 22.02 ± 9.67 % for loaded-POs compared to 53.50 ± 10.82 % free. Notably, with a 1000-fold reduction in drug concentration, PXT-loaded POs exhibited a toxicity equal to that reached by the free drug, which was maintained for all the measurements taken throughout 24 hours (Fig.S6). Similarly, intracellular delivery of PNB using POs resulted in a 100-fold increase in the efficacy of PNB compared to the free drug. Lastly, CRF exhibited no effect on SF8628 cells either encapasulated or free during the initial 6 hours of treatment. At a late time-point, CRF and CRF-loaded POs showed similar cytotoxicity.

**Figure 2:**
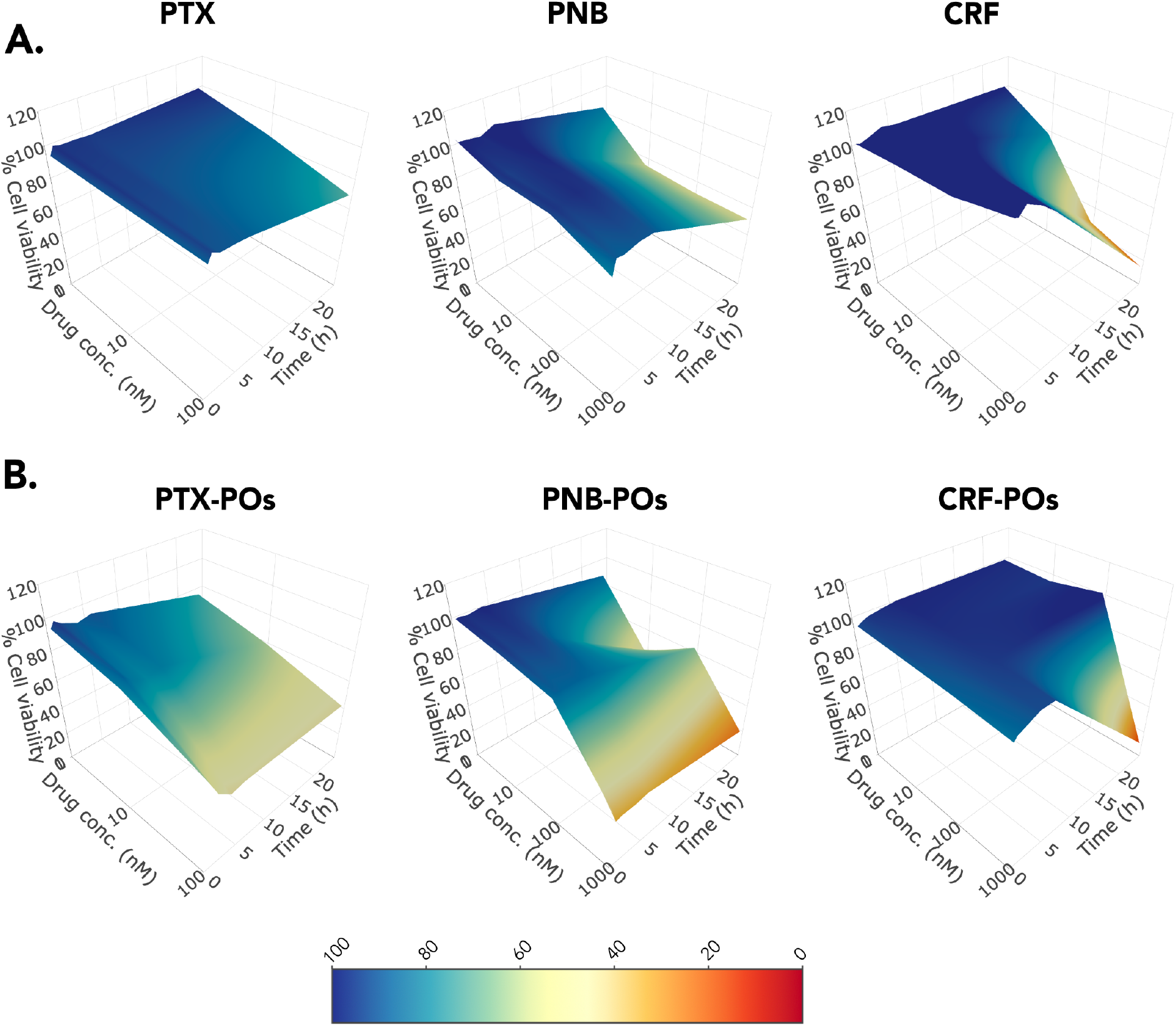
Effect of PXT, PNB and CRF-loaded PMPC-PDPA POs in cell viability of paediatric glioma cells. 3D-surface plots of cell viaiblity of SF8628 cells treated with either free PXT, PNB or CRF solubilised in DMSO (A) or PXT-POs, PNB-POs or CRF-POs (B) over 24 hours of incubation and different drug doses. The colour bar indicates percentage of cell viability.

In order to establish a panel of drug combinations, SF8628 cells were incubated with three different combinations: PNB:PXT, PNB:CRF and PXT:CRF, at different ratios. These experiments generated 120 data points of cell viability for each of the three combinations over 24 hours, which were plotted as heatmaps for an easier interpretation (Fig.3). Additionally, cell viability at 24 hours is shown separately (Fig.3A-C). Generally, drug combinations were more effective when entrapped within the POs, as a lower cell viability was obtained for the POs formulations. Particularly, combinations of PNB:PXT at ratios 100:100, 10:100 and 1:100 (nM), and of PXT:CRF at ratios 100:1, 100:10, 100:100 and 10:100 (nM), reduced cell viability to significantly lower values (P ≺ .05) when administered as drug-loaded POs compared to free drugs (Fig.3). These results indicate that intracellular delivery of drug combinations enhances the efficacy of treatment in paediatric glioma cells. In contrast, no significant difference was observed between free and loaded drugs for the rest of the tested ratios at 24 hours in these two combinations. Interestingly, drug combination efficacy is enhanced with the intracellular delivery of high ratios of PTX, which could be due to the fast action of PTX on molecular pathways and further sensitisation to the other drug. PNB:CRF combination revealed no significant difference between free and drug-loaded formulation at any tested ratios. Notably, a faster kinetics in cytotoxicity is observed for the three combinations when encapsulated within the PMPC-PDPA POs.

**Figure 3:**
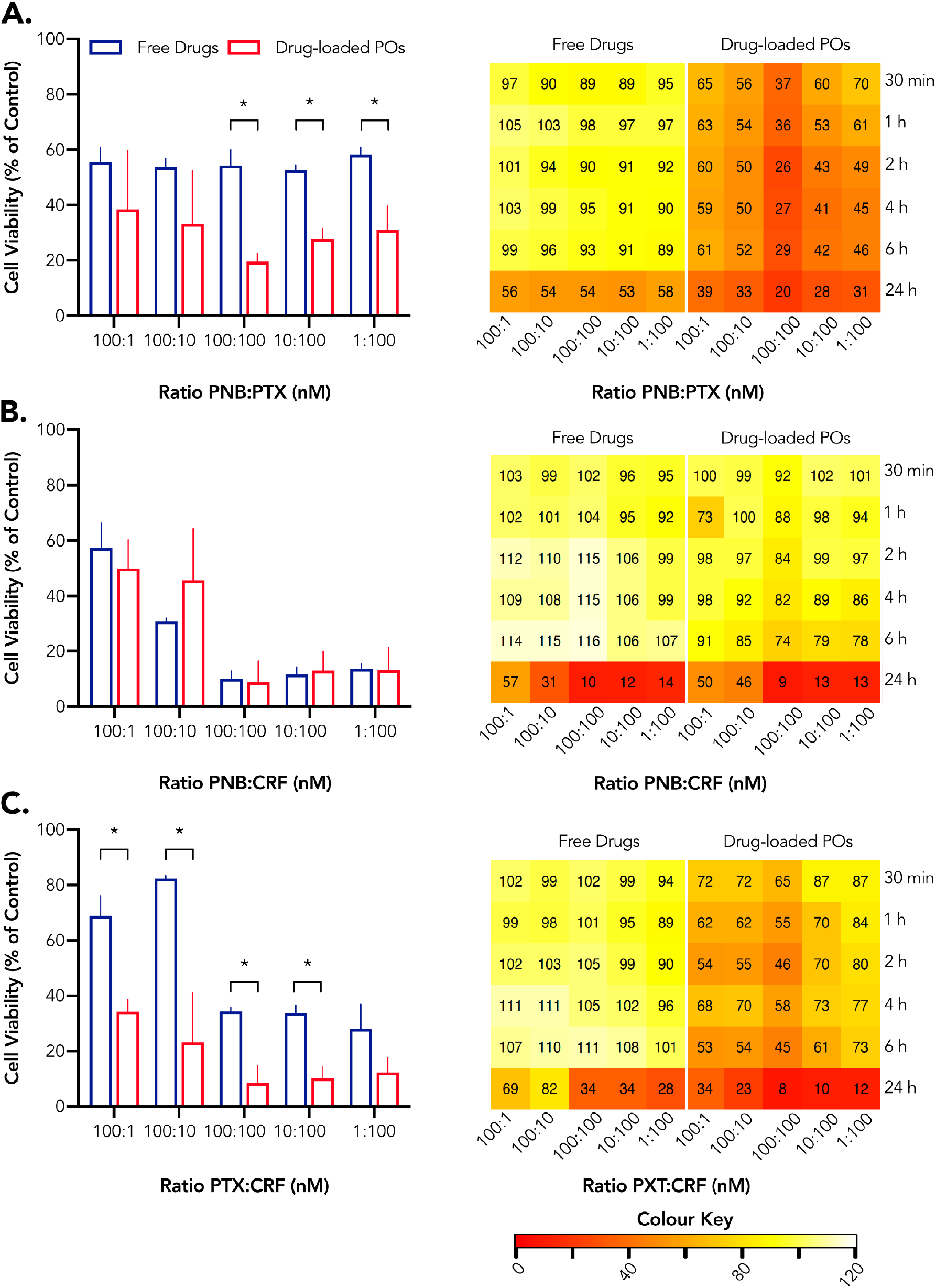
Effect of drug combinations in the cell viability of human SF8628 paediatric glioma cells. Cell viability of SF8628 treated with PNB:PXT (A), PNB:CRF (B) and PXT:CRF (C) after 24 hours of incubation together with heatmaps representing toxicity at 30 minutes, 1, 2, 4, 6 and 24 hours. The colour key indicates the relationship between colours and % of cell viability, where red indicates higher toxicity. Mean ± SD (n=2). * P<0.05, comparing free drugs to drug-loaded POs.

To asses whether intracellular delivery improves the synergistic effect of the three drugs, the pharmacological interaction of all tested combinations was evaluated using the R package “SynergyFinder”, which provides 4 synergistic scores for each combination using 4 different reference models: Loewe, Bliss, ZIP and HSA (S.8). Synergy score values > 0 were referred as synergistic, which means that the effect of each drug is enhanced, while synergy scores < 0 were denoted antagonistic, which indicates that the drugs interact in way that reduces the final therapeutic effect. As shown in Figure 4, intracellular delivery of drug combinations via pH-sensitive PMPC-PDPA POs consistently triggered enhanced synergy in the three tested drug combinations. Interestingly, the average synergy value for the PNB:PXT combination (Fig.4) denoted antagonistic interactions when incubated with the free drugs (except for the HSA model although this was very close to zero), while PNB:PTX co-delivery showed synergistic interactions when encapsulated in POs. Although the combination of free CRF with PNB (PNB:CRF) and PXT (PXT:CRF) demonstrated synergy, the average scores for these combinations were higher when encapsulated in POs. These findings demonstrate that intracellular drug delivery improves synergistic drug interactions in paediatric glioma cells reflecting the clinical potential of PMPC-PDPA POs to provide strong synergistic drug profiles.

**Figure 4:**
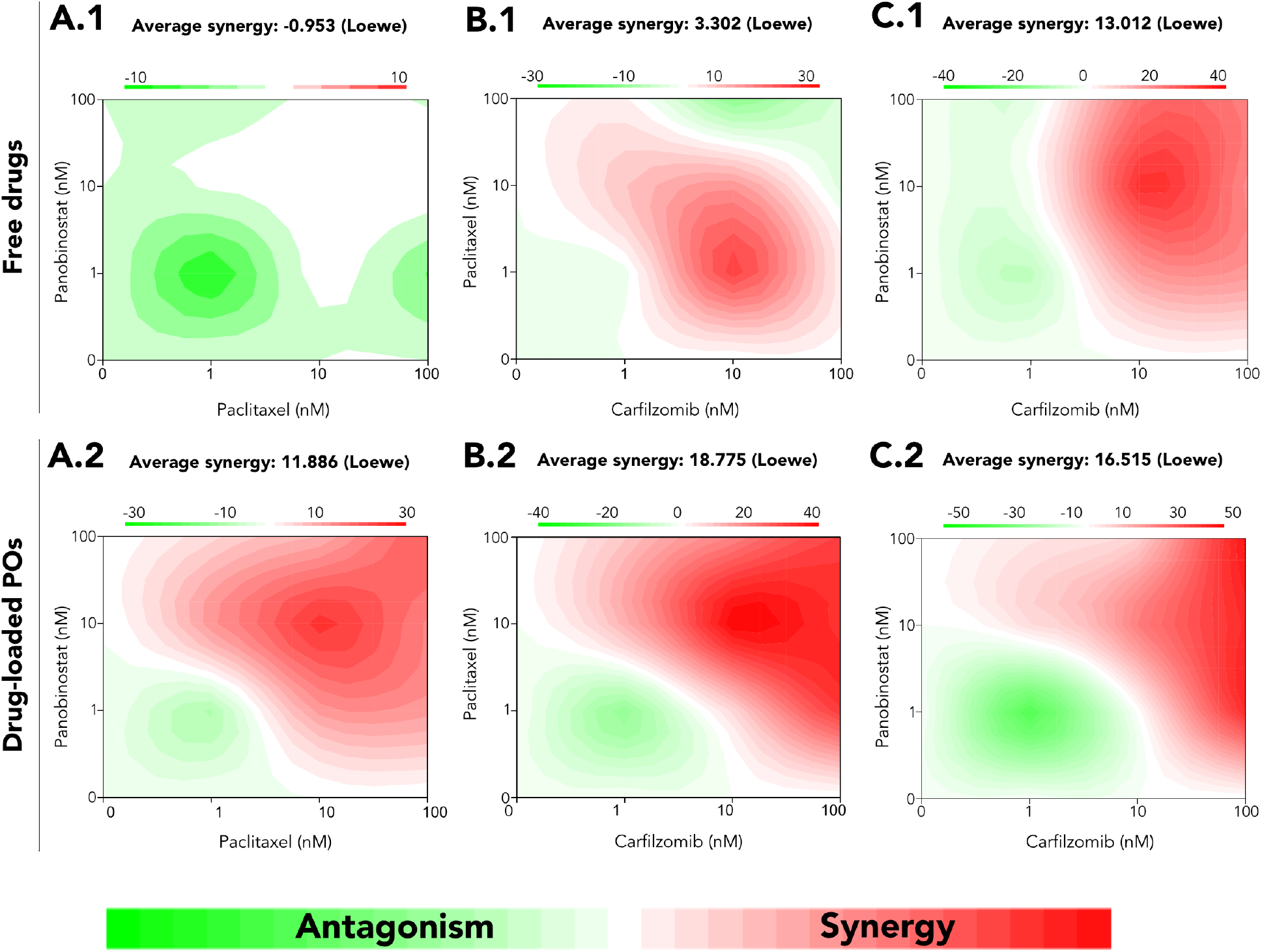
Drug synergy analysis. Dose–response matrix of PXT:PNB (A.1) and PXT-POs:PNB-POs (A.2) combinations, PNB:CRF (B.1) and PNB-POs:CRF-POs (B.2) combinations and PNB:PXT (C.1) and PNB-POs:PXT-POs (C.2)combinations on the paediatric human glioma cell line SF8628. Synergy values are determined using the LOEWE reference model using the R package SynergyFinder. Red values indicate synergistic combination, green values antagonistic interactions and white denote additivity.

In conclusion, we demonstrated that poorly water soluble drugs were successfully entrapped within PMPC-PDPA. Then, we further show that intracellular drug delivery mediated by pH-sensitive PMPC-PDPA POs improves the cytotoxcity of PTX, CRF and PNB either alone or in combination in paediatric glioma cells. Furthermore, intracellular delivery mediated by the POs leads to an enhanced synergistic effect of drug combinations (Fig.4 and S7). Most importantly, we show that using POs to solubilise poorly water-soluble drugs, the intracellular delivery of drug combinations by PMPC-PDPA POs allow us to obtain potent synergistic effects of anti-cancer drugs in glioma, which could not be identified using conventional drug solubilisation. This methodology can be applied to other panels of combined drugs in order to optimise the identification of the real synergistic potential of drugs combinations.

## Acknowledgements

G.B. thanks the ERC for the starting grant (MEViC 278793) and consolidator award (CheSSTaG 769798), the EPSRC/BTG Healthcare Partnership (EP/I001697/1), EPSRC Established Career Fellowship (EP/N026322/1) and EPSRC/SomaNautix Healthcare Partnership EP/R024723/1. G.B. and B.S thank Children with Cancer UK for the research project (16-227).

## Graphical TOC Entry

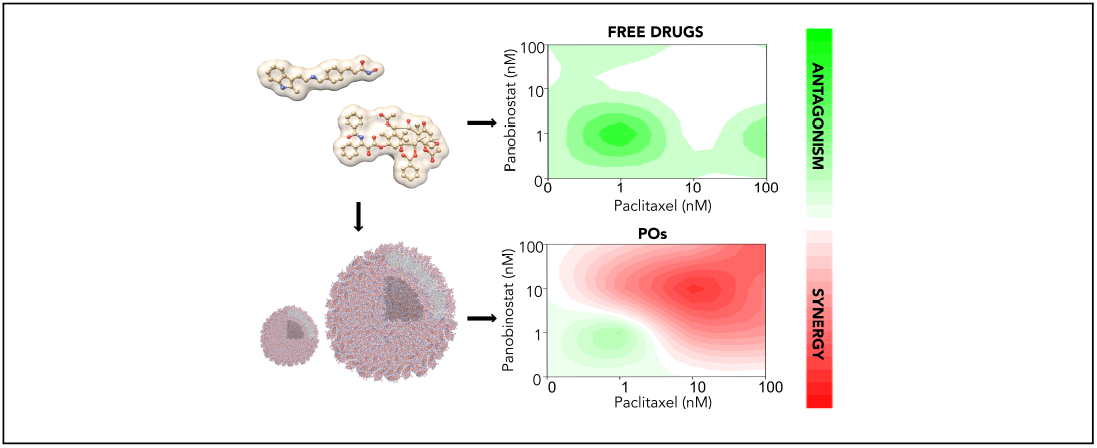

