## Supporting information for "Combinatorial Intracellular Delivery Screening (CIDS) of Anticancer Drugs"

**Abstract**

Conventional drug solubilisation strategies limit the understanding of the full potential of poorly water-soluble drugs during drug screening. Here, we propose a screening approach in which poorly water-soluble drugs are entrapped in poly (2-(methacryloyloxyethyl phosphorylcholine)-poly(2-(diisopropylaminoethyl methacryate) (PMPC-PDPA) polymersomes (POs) to enhance drug solubility and facilitate intracellular delivery. By using a human paediatric glioma cell model, we demonstrated that PMPC-PDPA POs mediated intracellular delivery of cytotoxic and epigenetic drugs by receptor-mediated endocytosis. Additionally, when delivered in combination, drug-loaded PMPC-PDPA POs triggered both an enhanced drug efficacy and synergy compared to that of a conventional combinatorial screening. Hence, our comprehensive synergy analysis illustrates that our screening methodology, in which PMPC-PDPA POs are used for intracellular co-delivery of drugs, allows us to identify potent synergistic profiles of anticancer drugs.

### Supplementary information

#### Characterisation of PMPC-PDPA POs by TEM

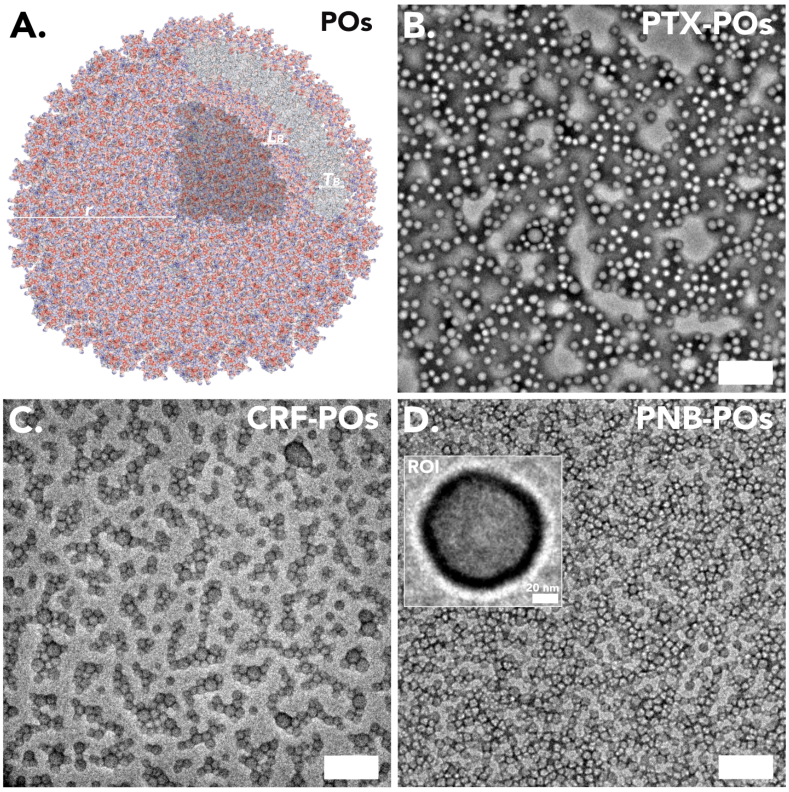


Figure S1: Morphology of PMPC-PDPA POs. (A) Schematic representation of the PMPC-PDPA PO structure showing the hydrophobic, $T_B$, and hydrophilic layers, $L_B$, and the vesicle radius, r. TEM images of (B) PTX-, (C) CRF- and (D) PNB-loaded POs showing monodisperse formulations with a size of ≈ 70 nm. The inset illustrates a region of interest (ROI) of a single PO with a membrane of dimensions of ≈ 6 nm. Scale bar: 300 nm.

#### Characterisation of PMPC-PDPA POs by HPLC

| | Size (nm) | PDI | Drug conc. ($\mu$M) | POs P.E. (%) | L.E. (n/psome) | Log P (%) |
|---|---|---|---|---|---|---|
| POs | 71.15 | 0.177 | - | 94.51 | - | - |
| CRF-POs | 74.8 ± 3.5 | 0.175 | 109.7 | 20.23 | 1.75E+04 | 3.1 to 4.2 |
| PXT-POs | 71.2 ± 3.3 | 0.166 | 3.4 | 5.9 | 58 | 3.2 to 3.5 |
| PNB-POs | 117.0 ± 7.0 | 0.466 | 15.4 | 30.25 | 120 | 1.5 to 3.2 |

Table S1. Characterisation of PTX-, CRF- and PNB-loaded PMPC-PDPA POs. Size (nm), PDI, drug concentration ($\mu$M), POs production efficiency (%) and loading efficiency (n/psome) of unloaded and drug-loaded PMPC-PDPA POs, together with the drug logP.

### Characterisation of PMPC-PDPA POs by DLS

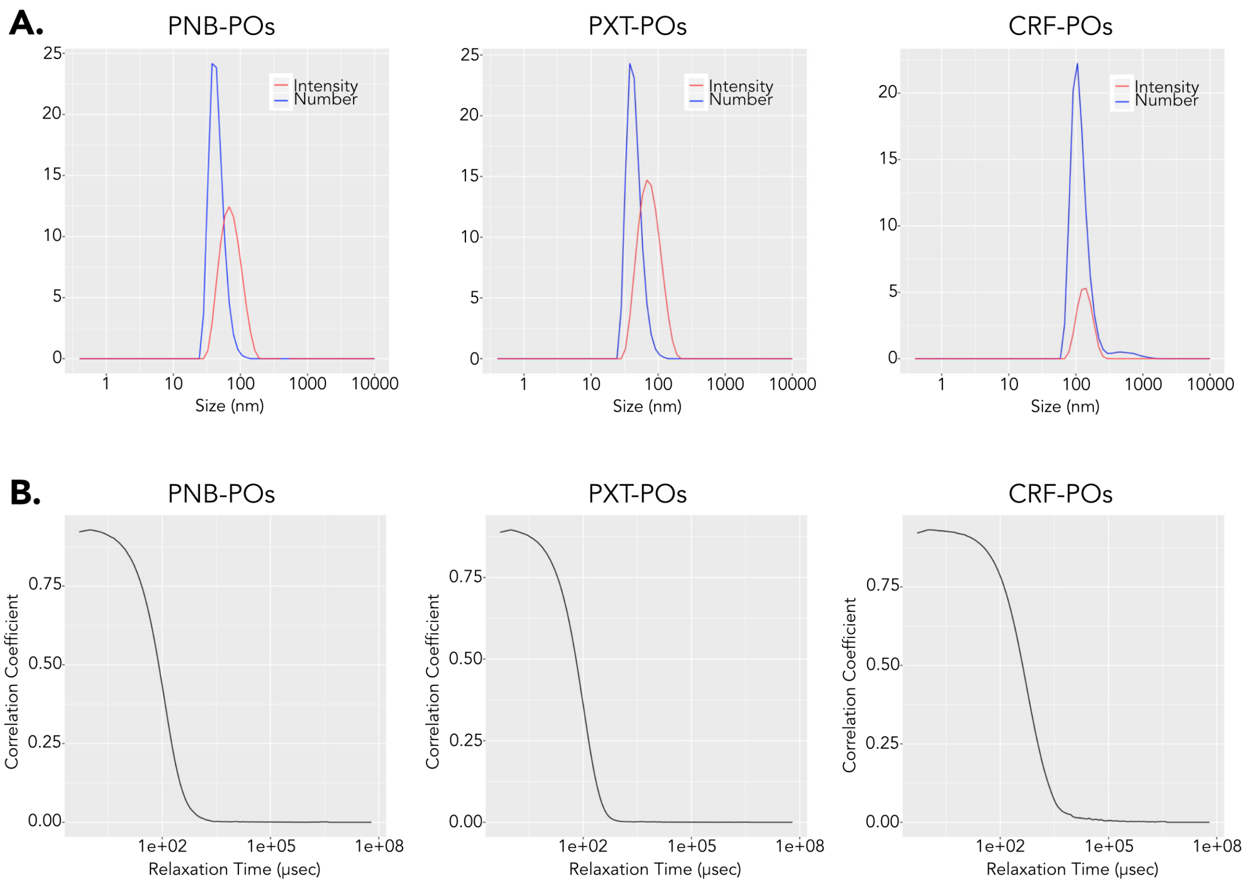


Figure S2: Characterisation of PMPC-PDPA POs by DLS. Histogram of PNB-, PTX- and CRF-loaded POs showing the size distributions expressed as % number (blue) and % intensity (red) and correlograms from POs samples produced by solvent switch.

#### PMPC-PDPA POs uptake by F98 cells

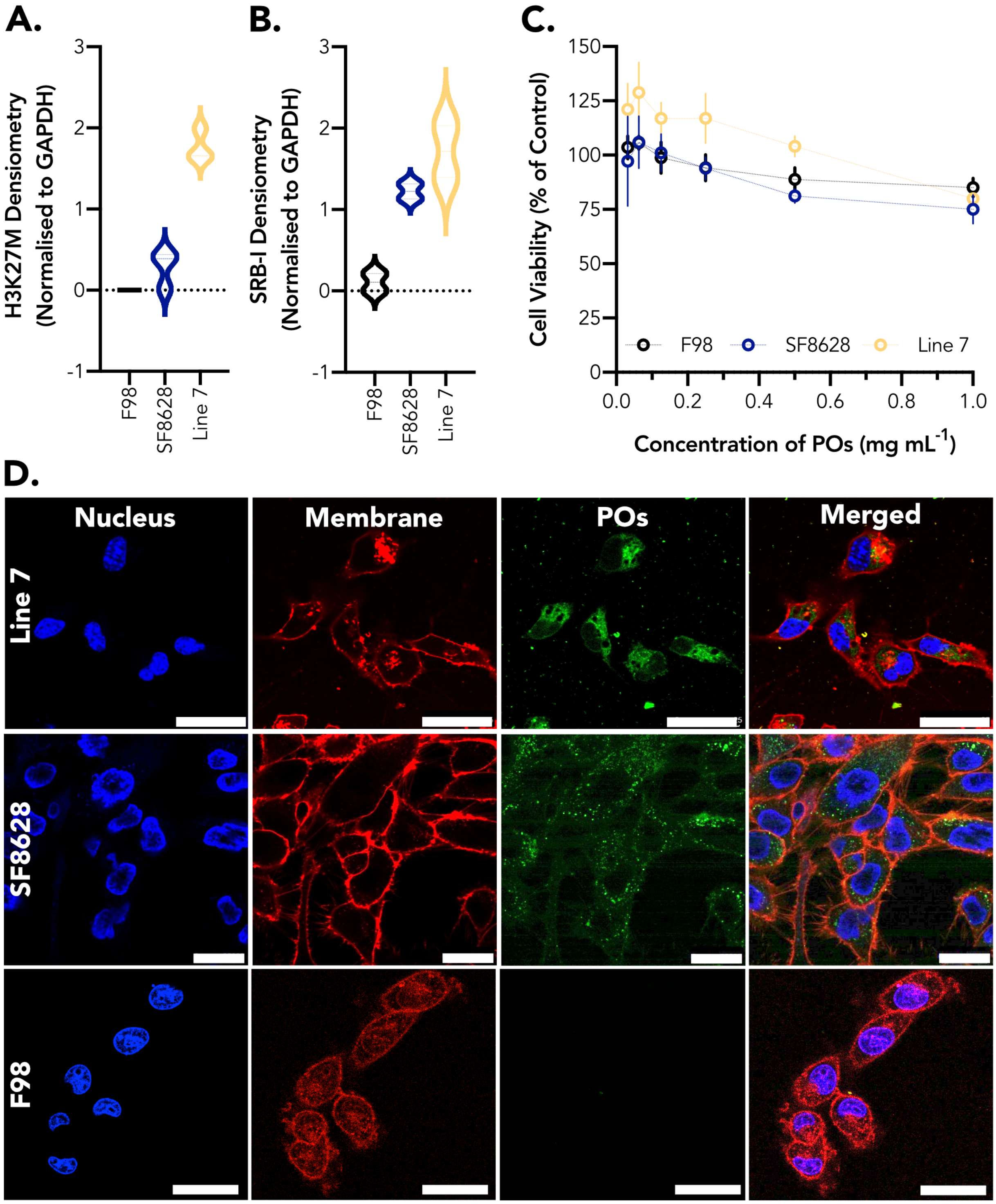


Figure S3: **PMPC-PDPA POs uptake by glioma cells**. Western blot analysis for the expression of (A) H3K27M mutation and (B) SRB1 receptors in glioma cell lines: cell line 7, SF8628 and F98; **(C)** MTT assay of empty PMPC-PDPA POs in glioma cells line SF8628. Mean ± SD (n=2g); (D) Confocal imaging of Rho-PMPC-PDPA POs uptake by the glioma cell lines. Blue: cell nucleus. Red: cell membrane. Green: fluorescent Rho-PMPC-PDPA POs. Scale bar = 25 $\mu$m.

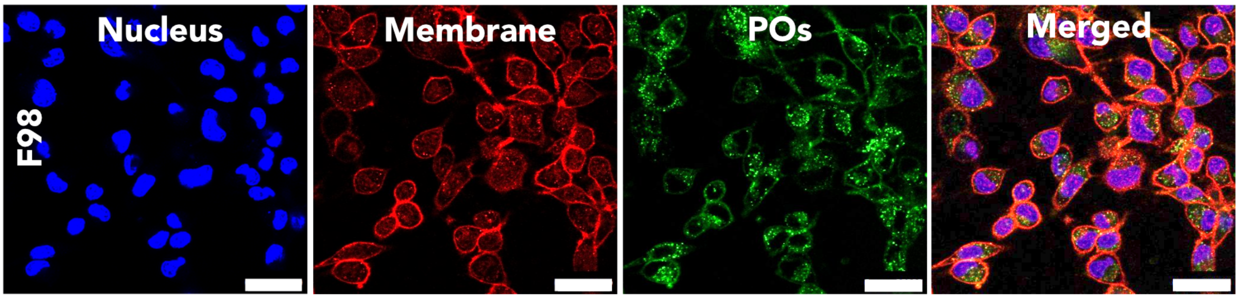


Figure S4: PMPC-PDPA POs uptake by F98 cells at 24 hours of incubation. Confocal imaging of Rho-PMPC-PDPA POs uptake by F98. Blue: cell nucleus. Red: cell membrane. Green: fluorescent Rho-PMPC-PDPA POs. Scale bar = 25 $\mu$m.

#### Cytotoxicity of PXT, CRF and PNB loaded PMPC-PDPA POs in paediatric glioma cells

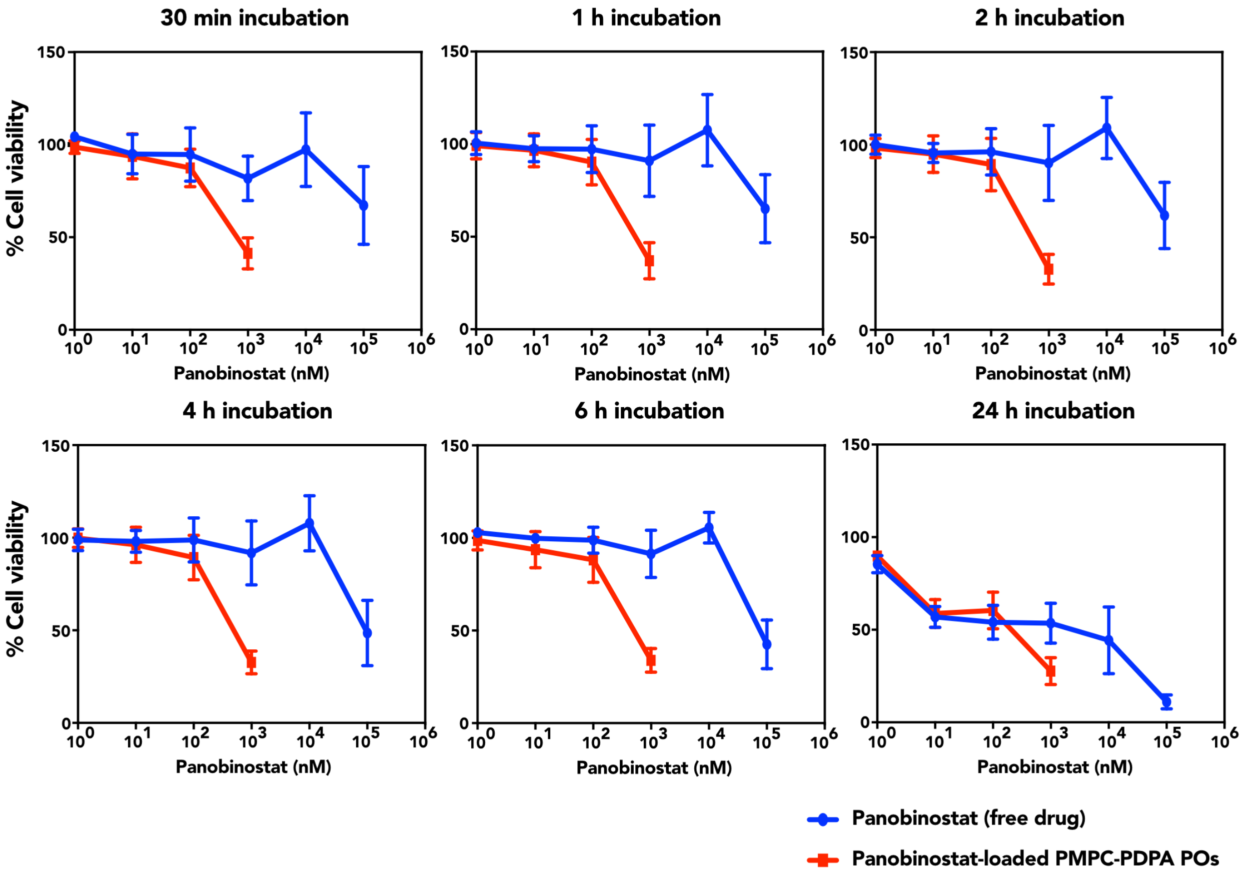


Figure S5: Dose-response curves of cell viaiblity of SF8628 cells treated with either free PNB dissolved in DMSO and PNB-POs over 24 hours of incubation and different drug doses. Mean ± SD (n=3).

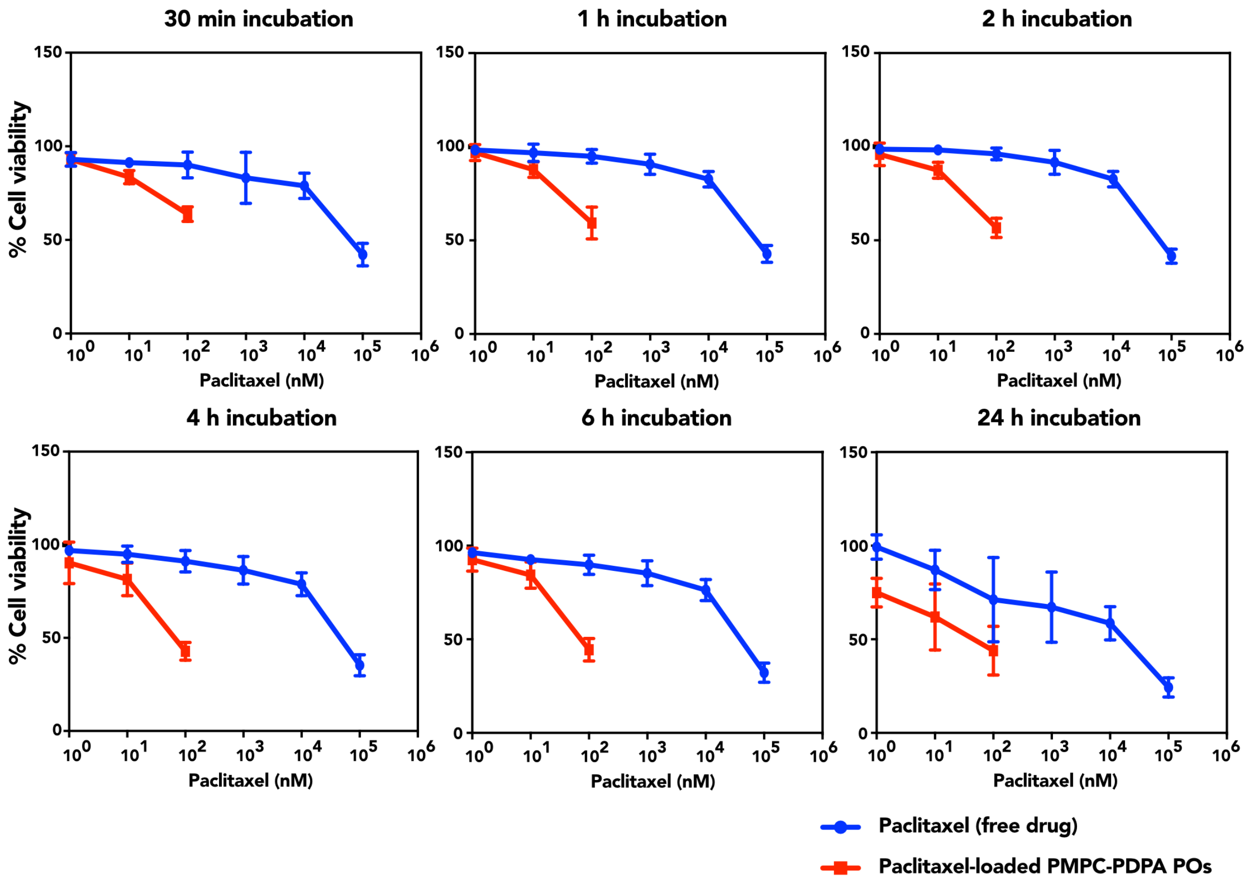


Figure S6: Dose-response curves of cell viaiblity of SF8628 cells treated with either free PXT dissolved in DMSO and PXT-POs over 24 hours of incubation and different drug doses. Mean ± SD (n=3).

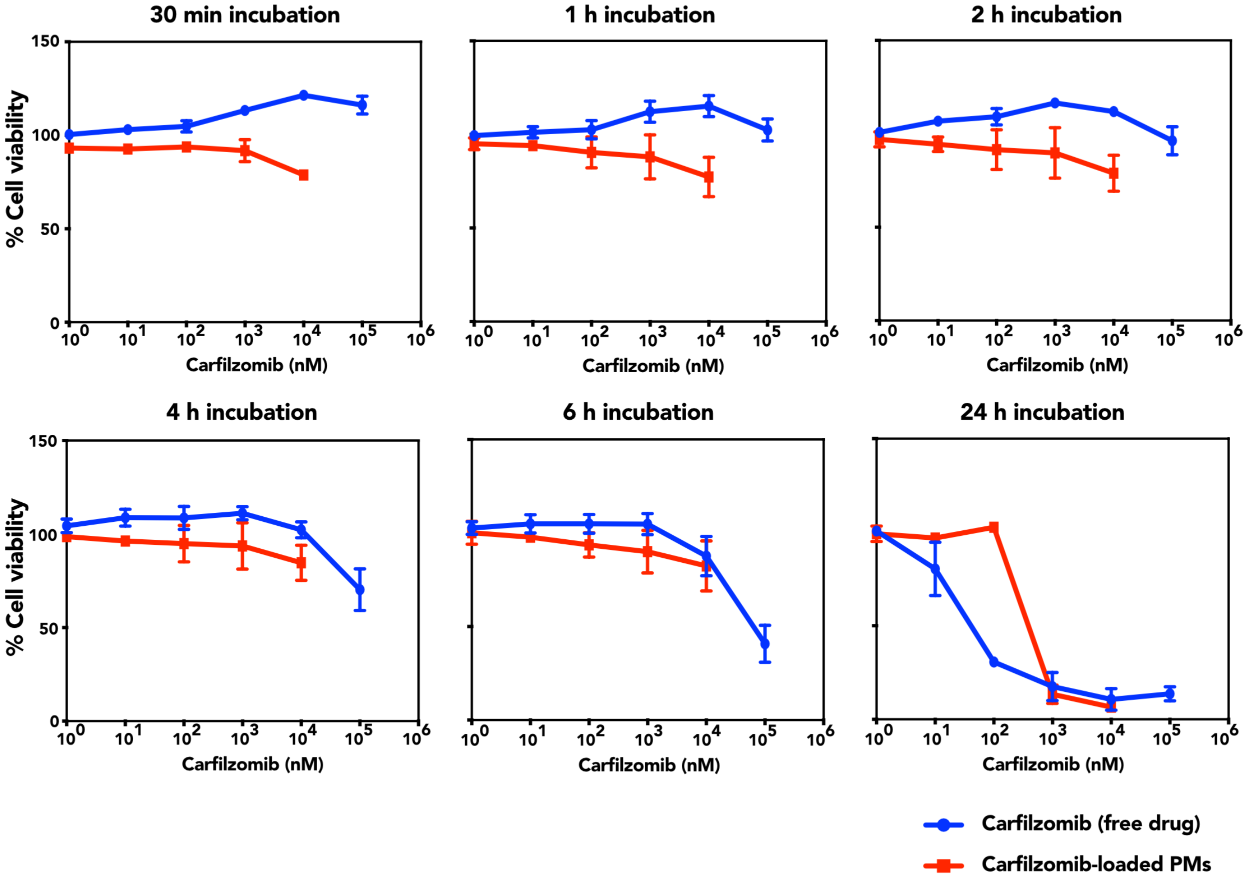


Figure S7: Dose-response curves of cell viaiblity of SF8628 cells treated with either free CRF dissolved in DMSO and CRF-POs over 24 hours of incubation and different drug doses. Mean ± SD (n=3).

***In vitro* screening of drug combinations on paediatric glioma cells**

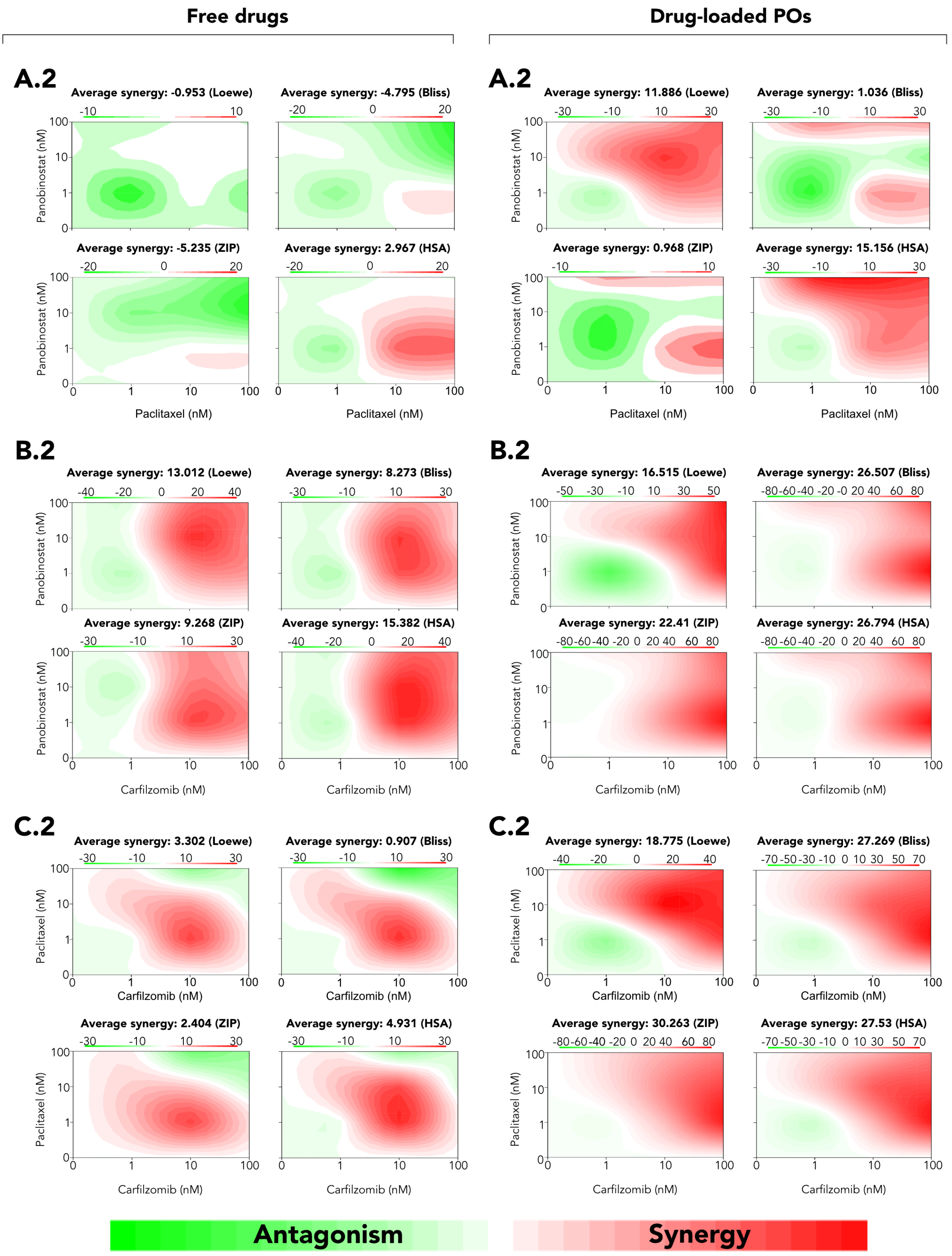


Figure S8: Drug synergy analysis. Dose–response matrix of PNB:PXT (A.1) and PNB-POs:PXT-POs (A.2) combinations, PNB:CRF (B.1) and PNB-POs:CRF-POs (B.2) combinations and PNB:PXT (C.1) and PNB-POs:PXT-POs (C.2)combinations on the paediatric human glioma cell line SF8628. Synergy values are determined using the LOEWE, Bliss, ZIP and HSA reference models using the R package SynergyFinder. Red values indicate synergistic combination, green values antagonistic interactions and white denote additivity.

### Materials and methods

**Materials** All reagents were obtained from Sigma-Aldrich unless stated otherwise. Drugs PTX, CRF and PNB were purchased from MedKoo Biosciences. Methanol, tetrahydrofuran and Spectra/Por dialysis tubing (3.5 kDa molecular weight cut off, 45 mm at-width) were obtained from Fisher Scientific. Phosphate buffered saline (PBS, pH 7.4), Dulbecco's Modified Eagle's Medium (DMEM), CellMask$^{TM}$ Deep Red Plasma Membrane Stain, Hoechst 33342 were obtained from Thermo Fisher Scientific. NeuroCult Proliferation Kit (Mouse & Rat) was purchased from StemCell Technologies. RealTime-Glo MT Cell Viability Assay was purchased from Promega. Protease inhibitors, BCA protein assay kit, and Laemmli sample buffer (x4) were purchased from Biorad. Rabbit monoclonal to H3K27M (ab190631) was obtained from Abcam. Rabbit polyclonal anti-SR-BI (NB400-104) was purchased from Novus Biologicals. SF8628 human DIPG H3.3.-K27M cell line was kindly provided by Prof. Rintaro Hashizume, Northwestern University. F98 undifferentiated malignant glioma cell line was purchased from ATCC. Cell line 7 was kindly provided by Prof. Paolo Salomoni, University College London.

**Polymersome preparation** $PMPC_{25}-PDPA_{70}$ and rhodamine B-$PMPC_{25}-PDPA_{70}$ polymers were synthesised by atom-transfer radical polymerisation (ATRP) synthesis as previously reported ([?] ). Self-assembly of $PMPC_{25}$-$PDPA_{70}$ POs was obtained by using a solvent switch method.[?] Briefly, $PMPC_{25} - PDPA_{70}$ (20 mg) was dissolved in 3:1 (v/v) methanol:tetrahydrofuran to a final concentration of 10 mg/ml and then, phosphate buffered saline (PBS, pH 7.4, 2.3 ml) was added at a flow rate of 1 $\mu$l per min under mechanical stirring at 40 °C. Fluorescently labelled POs were prepared by adding rhodamine B-$PMPC_{25} - PDPA_{70}$ at 1 mg/ml to the organic solution of PMPC-PDPA. PXT, CRF and PNB loaded $PMPC_{25} - PDPA_{70}$ POs were also prepared by solvent switch method by dissolving the drug in the organic solution containing PMPC-PDPA. To help drug solubilisation, solutions were sonicated and heated up to 40 °C for 10 minutes when necessary. After preparation, all formulations of PMPC-PDPA POs were purified by dialysis (3.5 KDa cut-off) against PBS (pH 7.4) followed by size exclusion chromatography (45-165 $\mu$m bead diameter), as previously reported.[?]

**Transmission electron microscopy.** The morphology of the PMPC-PDPA POs was evaluated using transmission electron microscopy (TEM). Briefly, 5 $\mu$L of PO sample was placed on a glow-discharged copper grid for 1 minute. Then, the sample was removed by absorption with a filter paper and the grid was immersed into a 20 $\mu$L drop of phosphotungstic acid (PTA) for three seconds. PTA was quickly removed and the grid was dried under vaccum for 20 seconds. Images were acquired on a FEI Tecnai G2 Spirit TEM microscope at 80 kV.

**Dynamic light scattering.** Hydrodynamic diameter and polydispersity (PDI) were determined by dynamic light scattering (DLS) using a Zetasizer Nano ZS (Malvern Instruments) with a 120-mW He-Ne Laser at 630 nm in a scattering angle of 173 °. Formulations were diluted in filtered PBS (pH 7.4) to a concentration of 0.2 mg $mL^{-1}$ and analysed in a polystrene cuvette at 25 °C. Data was processed using a Dispersion Technology Software (Malvern Instruments).

**High performance liquid chromatography.** The amount of polymer and drug in each prepared formulation were quantified by reverse-phase high-performance liquid chromatography (RP-HPLC) using a C18 reverse-phase Jupiter column Phenomenex™ and a Dionex UltiMate 3000 instrument (Thermo Fisher Scientific). Both POs and drugs were analysed at 220 nm. The analysis was carried out using an elution gradient method consisting of two different mobile phases: 0.05% trifluoroacetic acid (TFA) in methanol (phase A) and 0.05% TFA in water (phase B). The separation method used is described as follows: 5% A to 100% A in 10 minutes following a linear gradient, then 100% A is maintained for 15 minutes and finally returned to 5% A in one minute in a lineal gradient, which is maintained for 6 minutes. HPLC data was analysed with Thermo Scientific Chromeleon Chromatography Data System (CDS) software. The formation of both empty and drug-loaded POs were characterised with the parameter of "Polymersomes Production Efficiency", defined as the percentage of initial mass of polymer used in the process of POs preparation that self-assembles into POs, and is calculated as:

$$\text{POs production efficiency (\%)} = \frac{\text{Final mass of polymer}}{\text{Initial mass of polymer}} \times 100 \quad (1)$$

The drug encapsulation process was characterised with the parameter of "Loading Efficiency", which was defined as the number of drug molecules loaded in each polymersome, using a previously reported method.[?]

**Cell culture** Human paediatric H3K27M cells (SF8628) were used between passage 5 and 14. Undifferentiated glioma cells F98 were used between passage 4 and 15. Mouse H3K27M Line 7 were used between passage 4 and 15. SF8628 and F98 were maintained in DMEM supplemented with 10% (v/v) FBS and 1% (v/v) P/S . Line 7 was maintained in NeuroCult Proliferation media. Cells were maintained at 37 °C in a humidified atmosphere of 5% $CO_2$ and the medium refreshed every 2-3 days. For subculture, cells were washed twice with PBS, incubated with 0.25% trypsin-EDTA for 3 minutes, centrifuged and resuspended in fresh media. Media was changed every 2-3 days.

**Western blot.** Cell samples were washed twice with PBS and then, RIPA buffer containing protease inhibitors (1:50) was added to the samples and left on ice for 1 hour. Cells were collected, centrifuged and the supernatant was collected for Western Blot analysis. Protein levels in the cell lysates were determined using BCA Protein Assay Kit. Proteins (10 $\mu$g) were separated on 10% SDS polyacrylamide gels and transferred to polyvinylidene difluoride (PVDF) membranes. Membranes were blocked with 5% (w/v) non-fat milk in Tris-buffered saline (TBS) containing 0.1% (w/v) Tween-20 (TBS-T) for 1 hour and then incubated with the correspondent antibody overnight at 4 °C. After washing with TBS-T, the membranes were incubated with a secondary antibody for 2 hours at room temperature and imaged using an Odyssey CLx (LI-COR Biosciences). The membranes were further probed for GAPDH as a loading control.

**Cellular uptake.** Cells were seeded on a 35 mm imaging dish (Ibidi™) at a density of 5 x $10^4$ cells per well and cultured overnight. For the imaging, cell membrane was stained with CellMask™ Deep Red Plasma membrane Stain (1:2000 dilution with PBS) for 8 minutes and cell nuclei with Hoechst Stain solution (1:2000 dilution with PBS) for 5 minutes. Cells were incubated with fluorescently tagged Rho-$PMPC_{25} - PDPA_{70}$ POs diluted in cell medium (1 mg $mL^{-1}$) and live imaging was performed at 37 °C to study the POs internalisation at different time-points. Images were acquired using a Leica TCS SP8 confocal microscope equipped with Diode 405, Argon, DPSS 561 and HeNe633 lasers. Images were acquired via sequential scan to reduce fluorophore bleed-through. For live-cell imaging, an incubator at 37 °C and 5% $CO_2$ connected to the unit was used, and allowed to stabilise for 1 hour before imaging. Images were acquired with a 63x oil immersion objective at 400Hz and 512x512 pixels. Data analysis of acquired images was carried out using with ImageJ software. Cellular uptake was also analysed by flow cytometry. Briefly, after incubation with Rho-$PMPC_{25} - PDPA_{70}$ POs, cells were fixed using 4%(v/v) paraformaldehyde (PFA). Flow cytometry was performed using BD LSRFortessa™ Cell Analyzer (BD Biosciences). Untreated cells were used as a negative control to subtract autofluorescence. Data analysis was carried out using FlowJo software.

**Cell viability.** Cell viability was assessed by using the RealTime-Glo MT Cell Viability Assay. Cells were plated in a 96 well plate at a density of 8 x $10^3$ cells per well and then treated with increasing concentrations of free drugs or drug-loaded POs ranging from 1 nM to 100 $\mu$M. RealTime-Glo™MT Cell Viability Assay was used as specified by the supplier and allowed for a continuous measurement of cell viability up to 72 hours using a plate reader luminometer (Spark Multimode microplate reader, Tecan). The obtained data was analysed with Graphpad Prism™.

**Synergy Analysis.** Synergy of tested drug pairs was evaluated with the R package SynergyFInder (from within R, enter citation("synergyfinder")). SynergyFinder calculates the synergy scores (SS) and generates 2D plots of the synergy. The SS is calculated by comparing the experimental effect of a drug combination with their theoretical "expected" effect, where the latter is obtained based on the effect of the individual drugs either free or loaded, accordingly. If the experimental effect is higher than the expected one then synergy is claimed. Conversely, a measured effect lower than the theoretical would mean antagonism. Finally, the combination is called additive if both effects are the same. As up-to-date there is not much consensus on which reference model is the best to determine the expected effect, synergy was examined here by four different null models: Loewe, Bliss, HSA and ZIP. The difference between the four methods relies on the way in which the expected effect has been calculated.The R-package and its source-code are freely available at https://bioconductor.org/packages/ release/bioc/html/synergyfinder.html .

**Statistical Analysis** The results are expressed as mean ± standard deviation (SD). Statistical differences were evaluated by Student's t test in GraphPad Prism 7.03, and $P < .05$ was considered as statistical significant.
